# Evaluation of MALDI-ToF Mass Spectrometry for Rapid Detection of Cereulide from *Bacillus cereus* Cultures

**DOI:** 10.1101/869958

**Authors:** Joerg Doellinger, Andy Schneider, Timo Stark, Monika Ehling-Schulz, Peter Lasch

**Author notes:** Correspondence to: Joerg Doellinger.

## Abstract

*Bacillus cereus* plays an often unrecognized role in food borne diseases. Food poisoning caused by this pathogen is manifested by either diarrhea or emesis. While different enterotoxins have been linked to the diarrheal type of *B. cereus* infections, the emetic toxin cereulide is responsible for the second type. Due to the relatively high prevalence of cereulide associated food poisoning, methods for simple and reliable detection of cereulide producing strains are of utmost importance. Currently, liquid-chromatography coupled to mass spectrometry (LC-MS) is used for sensitive, specific and quantitative cereulide detection, but this technique requires specialized LC-MS equipment, which is often not available in microbiology routine diagnostic laboratories.

The last decade has witnessed the advent of matrix-assisted laser desorption/ionization-time-of-flight mass spectrometry (MALDI-ToF MS) as a simple, rapid and cost-efficient technique for identification of microbial pathogens in routine diagnostics. Just recently, two different studies reported on the application of MALDI-ToF MS for either the differentiation of emetic and non-emetic strains of *B. cereus* or for direct detection of cereulide from bacterial colony smears. However, no method evaluation and optimization was performed in frame of these studies. Thus, additional investigations on the selectivity and sensitivity of MALDI-TOF MS for cereulide detection are needed before implementation of this method in routine diagnostics can be considered. These aspects prompted us to investigate open or controversial issues and to systematically test sample preparation methods, commonly used for microbial identification for their suitability to detect the emetic toxin directly from bacteria.

Based on our experimental findings we propose a MALDI-ToF MS workflow that allows identification of *B. cereus* and sensitive detection of cereulide in parallel, using standard, linear-mode MALDI-ToF MS equipment. The experimental protocol is based on the well-established ethanol/formic acid extraction method and offers, if required, possibilities for further characterization by more sophisticated LC-MS-based methods. In summary, the ease of use and the achieved level of analytical sensitivity as well as the wide-spread availability of standard MALDI-ToF MS equipment in clinical microbiological laboratories provides a promising tool to improve and to facilitate routine diagnostics of *B. cereus* associated food intoxications.

## 2. Introduction

*Bacillus cereus* is a facultative anaerobic member of the Genus *Bacillus*, which is ubiquitously distributed in nature and commonly isolated from soil and food. It is a well-recognized causative agent of two different types of gastrointestinal diseases mediated by toxins but is also increasingly reported to be linked to non-gastrointestinal diseases, such as nosocomial or eye infections (Messelhäußer and Ehling-Schulz, 2018;Ehling-Schulz et al., 2019). The protein complexes non-hemolytic enterotoxin (Nhe) and hemolysin enterotoxin BL (Hbl), as well as cytotoxin K1 (CytK1) are linked to diarrheal symptoms (Granum et al., 1993;Rowan and Anderson, 1998;Clavel et al., 2004), while the cyclic dodecadepsipeptide cereulide is the etiological agent of the emetic form of the foodborne disease (Agata et al., 1995). The diarrheal form is presumably caused by enterotoxin production of *B. cereus* in the intestine, resulting from the consumption of food contaminated with bacterial spores (Granum et al., 1993;Rowan and Anderson, 1998;Clavel et al., 2004). The emetic syndrome causing cereulide is, by contrast, preformed in foods contaminated with emetic strains of *B. cereus* prior to ingestion. Although emetic food poising has been primarily linked to starch-rich foods, such as rice, pastry and noodles, it is becoming increasingly evident that emetic strains can contaminate a much broader diversity of foods (Messelhausser et al., 2014;Rouzeau-Szynalski et al., 2020). Intoxication caused by cereulide leads to nausea and vomiting after 0.5 – 6 h incubation and lasts up to 24 h (Ehling-Schulz et al., 2004). The emetic symptoms are usually self-limiting, however severe cases, such as liver failure and acute encephalopathy, have been reported (Mahler et al., 1997;Ehling-Schulz et al., 2004;Fricker et al., 2007;Ichikawa et al., 2010). *In vitro* studies have shown that pancreatic beta cells are especially sensitive to the cereulide, resulting in cell death even at low concentrations (Virtanen et al., 2008;Vangoitsenhoven et al., 2014), and a potential link between emetic *B. cereus* and type-2 diabetes is currently under debate (Vangoitsenhoven et al., 2015). Using a pig model, it was shown recently that cereulide is taken up by the body, efficiently translocated and accumulate in certain host organs and tissues (Bauer et al., 2018).

The heat- and acid-stable cereulide is synthesized by a non-ribosomal peptide synthetase (NRPS), which is encoded by the cereulide synthetase (*ces*) gene (Ehling-Schulz et al., 2005b). The *ces* operon is located on a megaplasmid that shares its backbone with the pX01 toxin plasmid from *Bacillus anthracis* (Ehling-Schulz et al., 2006). Active production of the toxin is known to be stimulated by different external factors, such as temperature, type of nutrition and some others (Haggblom et al., 2002;Kranzler et al., 2016;Rouzeau-Szynalski et al., 2020). Cereulide acts as a cationic ionophore that inhibits mitochondrial activity (Mikkola et al., 1999). The cyclic dodecadepsipeptide cereulide has a molecular weight of 1152.5 Da and contains three repeats of the amino acids L-O-Val-L-Val-D-O-Leu-D-Ala. A number of variants of cereulide, such as isocereulides A-G (Marxen et al., 2015) or homocereulide (Pitchayawasin et al., 2004;Naka et al., 2019) have been described in detail, which vary in their ionophoric properties and toxicity, thereby demonstrating the remarkably high level of chemodiversity of this potent toxin.

*Bacillus cereus* can be frequently found as a microbiological contaminant in a wide range of different foods. As an endospore forming bacterium it is difficult to eliminate from the food chain and therefore is considered a major problem in food production processes (Andersson et al., 1995;Rouzeau-Szynalski et al., 2020). The prevalence of the bacteria in food samples related to food poisoning has been shown to be as high as 50 % (Messelhausser et al., 2014). Most of the *B. cereus* strains from randomly collected food samples are able to produce at least one of the diarrheal toxins (> 90 %) while only a small portion is able to produce the emetic toxin cereulide (1-10 %) (Hoton et al., 2009;Messelhausser et al., 2014). Although the prevalence of emetic strain is generally rather low, up to 30 % of certain food classes linked to foodborne outbreaks contained emetic *B. cereus* strains (Hoton et al., 2009;Messelhausser et al., 2014). It is thought that the incidence of foodborne *B. cereus* intoxications is currently underestimated, due to short duration of illness, missing awareness in the clinical field and the fact that reporting of *B. cereus* intoxications to authorities is obligatory in many countries only in case of outbreaks (EFSA, 2016). Furthermore, detection of the cereulide toxin itself is tedious and rather challenging due to the comparably low toxic dose and the lack of enrichment strategies.

A simple, reliable and sensitive direct detection method for cereulide in frame of routine microbiology diagnostics would therefore be desirable. Polymerase chain reaction (PCR) is used to screen bacterial isolates for the cereulide synthetase gene *ces*, which is however not a verification of the actual toxin production (Fricker et al., 2007) and recently a machine learning based Fourier Transform Infrared (FTIR) spectroscopy method for discrimination of emetic and non-emetic strains has been described (Bagcioglu et al., 2019). Currently, liquid-chromatography coupled to mass spectrometry (LC-MS) is the gold standard for sensitive cereulide detection and quantification (Bauer et al., 2010;Biesta-Peters et al., 2010;Stark et al., 2013). Based on a stable isotope dilution analysis (SIDA) assay (Bauer et al., 2010), an ISO method (EN-ISO, 18465) for accurate quantitation of cereulide in complex matrices, such as foods, has been established lately.

However, the latter technique is complex and is available mostly in specialized analytical laboratories. Within the last decade matrix-assisted laser desorption/ionization (MALDI) – time–of– flight mass spectrometry (MALDI-ToF MS) has revolutionized the way pathogens are identified. (Seng et al., 2009;Nomura, 2015;Schubert and Kostrzewa, 2017;Welker et al., 2019). Today, MALDI-ToF MS is used worldwide in laboratories for routine identification of clinically relevant pathogenic microorganisms. The technique offers a number of advantages over classical established methods, which has led not only to high level of acceptance of the technique but resulted also in wide dissemination of MALDI-ToF instrumentation in microbiological diagnostic laboratories.

Thus, it was evident that the potential of standard MALDI-ToF MS-based biotyping, carried out in the m/z range between 2 and 20 kDa, has been tested for differentiation of emetic from non-emetic *B. cereus* group members (Fiedoruk et al., 2016) and, more recently, also for direct detection of cereulide (Ducrest et al., 2019). The suitability of MALDI-ToF MS for differentiation between emetic and non-emetic *B. cereus* strains was shown by statistical analyses of the linear-mode low-mass MALDI-ToF MS data from 121 *B. cereus* group strains (Ulrich et al., 2019). A diagnostic accuracy of 99.1% was achieved. However, the specific biomarkers used for differentiation were found at m/z positions at 1171 and 1187 and would give an inexplicably large difference of approx. 4000 ppm, provided that the observed biomarkers represent the sodium and potassium adducts of the cereulide. As the molecular identity of the biomarker peaks was obviously not a subject of the aforementioned study, this inconsistency remains to be investigated. In another study, however, it has been demonstrated that reflectron-mode MALDI-ToF MS permits detection of the sodium and potassium adducts of cereulide at m/z 1175.6 ([M+Na]^+^) and m/z 1191.6 ([M+K]^+^), respectively (Ducrest et al., 2019). Cereulide was directly identified from colony smears of *B. cereus* while the limit of detection (LOD) of 30 ng/mL was established from a dilution series of standard cereulide. The same study proposed matrix-free detection by laser desorption/ionization (LDI) MS, which was reported to result in a 1000-fold improvement of LOD to 30 pg/mL (Ducrest et al., 2019). However, neither spectra nor calibration curves were provided that supported the claimed dramatic improvement of the analytic sensitivity for the emetic toxin.

Furthermore, both recently published studies (Ducrest et al., 2019;Ulrich et al., 2019) did not provide method evaluation or optimization of the experimental protocol for cereulide detection. The inconsistencies of the data and the limitations in the applicability of the proposed experimental protocols for simple, rapid and sensitive cereulide detection in a routine clinical laboratory have motivated us to systematically investigate the potential and limits of MALDI-ToF MS for cereulide detection in routine diagnostics.

To this end, we evaluated three different sample preparation methods commonly used in routine MS-based workflows for pathogen identification and tested different organic solvents for their ability to extract the emetic toxin from *B. cereus* materials. The experimental cereulide detection protocol was evaluated under blinded conditions and the LOD of cereulide detection from MALDI- and LDI-ToF MS measurements was determined. Furthermore, fragment analysis was performed for unambiguous identification of the emetic toxin by means of MALDI-ToF/ToF MS. Our study demonstrates that a routine sample preparation method for MALDI-ToF MS biotyping allows both, identification of *B. cereus* and parallel detection of the emetic toxin with sufficiently high analytical sensitivity. The proposed workflow, which can be carried out with routine linear mode MALDI-ToF MS instrumentation, represents a suitable high-throughput technique to screen *B. cereus* strains for their actual emetic toxin production.

## 3. Materials and Methods

### *B. cereus* strains

The *B. cereus* emetic reference strain F4810/72 (Ehling-Schulz et al., 2005a) was used for protocol optimization experiments, and a test panel of strains, derived from the *B. cereus* strain collection of the Institute of Microbiology at the Vetmeduni Vienna, was compiled for evaluation of the optimized protocol in frame of a blinded study (see table 1). The test panel, which comprised high, medium and low emetic toxin producing *B. cereus* strains as well as a so-called emetic-like (non cereulide producer, but genetically closely related to emetic strain (Ehling-Schulz et al., 2005a)), and the non-emetic *B. cereus* type strain was sent blinded by Vetmeduni to the Robert-Koch Institute (RKI) Berlin. Classification of strains as high, medium or low toxin producers was carried out using UPLC-TOF MS (Stark et al., 2013).

**Table 1:**
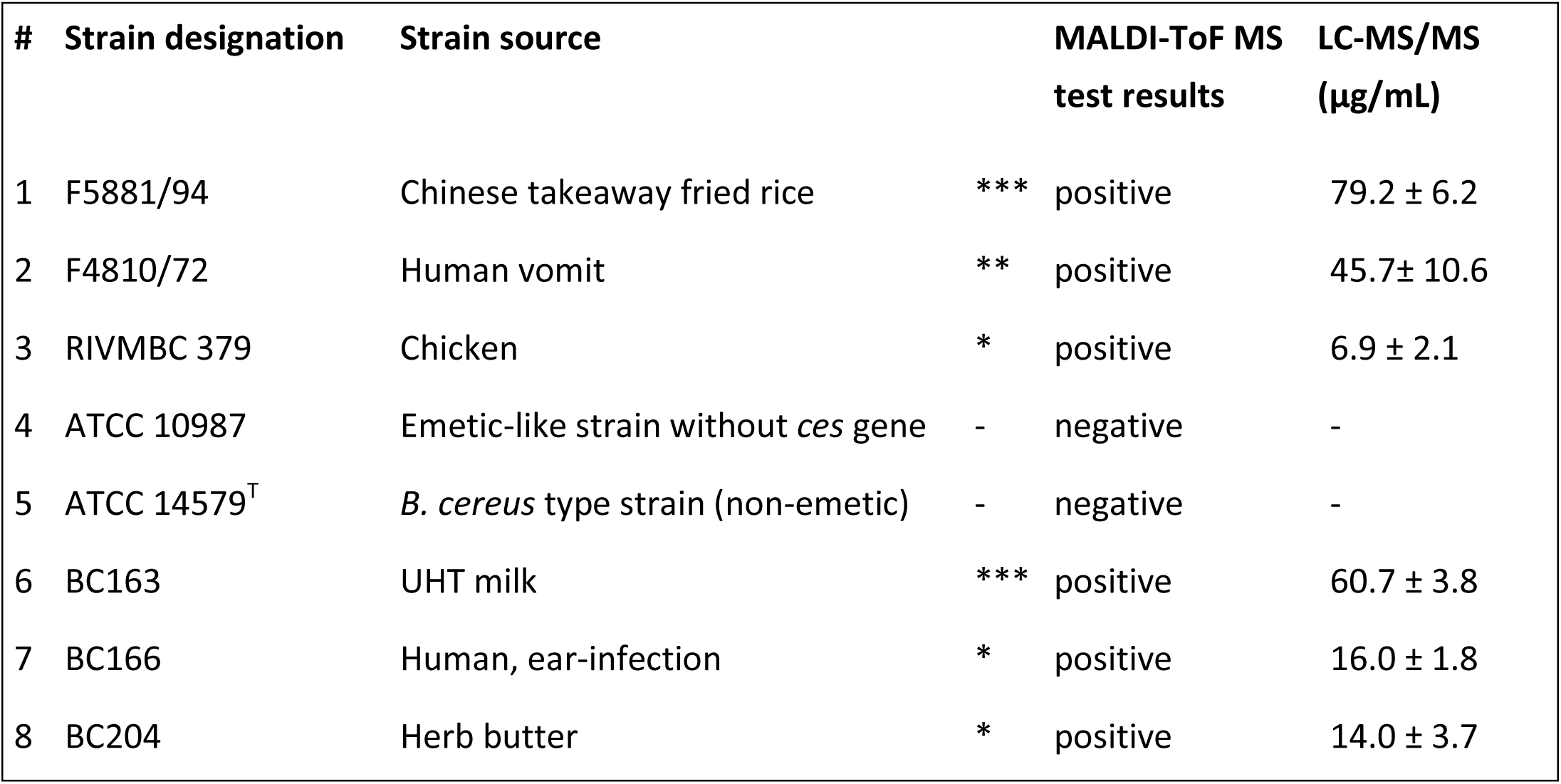
Analysis of cereulide in *B. cereus* strains by MALDI-ToF MS. Eight selected strains of *B. cereus*, differing in their cereulide production capabilities, were analyzed under blinded conditions using MALDI-ToF MS. For this purpose, the *B. cereus* strains were cultivated on Caso agar for 24 h at 37°C and *B. cereus* ethanol extracts were characterized by positive reflectron MALDI-ToF MS. Mass spectra exhibiting the sodium [M+Na]^+^, or potassium [M+K]^+^ adduct of cereulide at m/z 1175.6 and m/z 1191.6 with a SNR of at least 3 were regarded as cereulide positive. MALDI-ToF MS test results were compared after unblinding with the data derived from mass spectrometric profiling of *B. cereus* strains by LC-MS/MS. *** high-producer strain; ** medium-producer; * low-producer; - non-producer strain of cereulide; data according to (Stark et al., 2013)

### Cultivation of B. cereus for MALDI-ToF MS analysis

Bacteria were cultured on different media used in bacterial diagnostic labs under aerobic conditions for 24 h at 37°C before toxin extraction. The tested cultivation media included Casein-Soy-Peptone (Caso, Merck KGaA, Darmstadt, Germany), Luria-Bertani (LB, produced in house) or blood agar produced in house from Columbia blood agar base (CM0331, Oxoid, Wesel, Germany) and sheep blood (5%).

### Sample preparation for cereulide detection in bacterial biotyping MALDI-ToF MS workflows

Three sample preparation methods for bacterial biotyping by MALDI-ToF MS, (i) direct transfer, (ii) ethanol/formic acid (FA) and (iii) trifluoroacetic acid (TFA) extraction were compared for their suitability to detect the emetic toxin. Direct transfer of bacterial cells from blood, LB or Caso agar plates onto the MALDI target was performed according to the manufacturer’s instructions of the MALDI Biotyper system (MALDI Biotyper 2.0 User Manual, 2008, Revision 2, Bruker Daltonik, Bremen, Germany). The spots were overlaid with 1 μL of saturated matrix solution (12 mg/mL α-cyano-4-hydroxy-cinnamic acid, HCCA, in 70% acetonitrile, ACN, and 3.0% TFA) and allowed to dry.

TFA and ethanol/FA extractions of the different *B. cereus* cultures were carried out as previously published (Lasch et al., 2008;Freiwald and Sauer, 2009;Lasch et al., 2016). In case of TFA extraction 1 μL of the 1:10 diluted TFA extract was mixed on target with 1 μL of the saturated HCCA matrix solution. For ethanol/FA extraction *B. cereus* cells were brought in 300 μL double-distilled water before 900 μL ethanol were added; cf. (Freiwald and Sauer, 2009). Volumes of 1 μL of the 75% ethanol fractions were mixed on target with 1 μL of the above-mentioned saturated matrix solution. The remaining volume (∼ 1200 μL) of the ethanol solution was dried and resolubilized in a volume of 20 μL pure ACN. From the concentrated ACN extract a volume of 1 μL was spotted onto a ground steel target (Bruker) and mixed with 1 μL of saturated matrix solution before drying. 1 μL of the FA sample solution was dried on target, overlaid by 1 μL of the above-mentioned matrix solution and dried again. ACN extraction of the cereulide from *B. cereus* cells was carried out under identical conditions, except that water and ethanol were replaced by 1200 μL ACN.

### Cereulide extraction from B. cereus colony material by solvents

Colony material from blood cultures of *B. cereus* F4810/72 (24 h, 37°C) equivalent to three inoculation loops was suspended in 500 μL organic solvent (methanol, n-pentane, ethanol, ACN and acetone) by vortexing. The samples were sonicated for 5 min at room temperature in an ultra-sonication bath before insoluble material was removed by centrifugation for 5 min at 4000 g. The supernatants were evaporated to dryness in a vacuum concentrator (SpeedVac, Uniequip, Martinsried, Germany). Dried samples were resolubilized in 20 μL ACN; resolubilized samples are in the following termed concentrated extracts. For MALDI-ToF MS measurements, 1 μL of the concentrated extracts were mixed on target with an equal volume of saturated HCCA matrix solution (see above) and allowed to air-dry before MS measurements.

### Cereulide detection and characterization using MALDI-ToF and LIFT-ToF/ToF MS

Dried samples were analyzed using an *Autoflex Speed* MALDI-ToF/ToF mass spectrometer (Bruker) in positive ion reflectron mode. The instrument was controlled by the FlexControl Software (Version 3.3.108) supplied by Bruker. External calibration was carried out using the peptide calibration standard II (Bruker). Full scan mass spectra were acquired in the m/z range of 700 – 3500 at 1 kHz repetition rate of the Smartbeam II Nd-YAG laser (excitation wavelength λ=355 nm). Each recorded spectrum was summed from 8 profiles, which were accumulated from 1000 laser shots each. Peaks were picked after smoothing and baseline subtraction using the SNAP algorithm implemented in the FlexAnalysis software (v. 3.3.80, Bruker). Cereulide was further characterized by LIFT-ToF/ToF MS analysis (Suckau et al., 2003). For this purpose, fragmentation spectra were accumulated at 200 Hz laser repetition rate and elevated laser energies from 1000 shots in parent acquisition and 1000 shots in fragment acquisition mode. The precursor ion selection range was 0.85 % of the precursor mass.

Linear MALDI-ToF spectra were collected in the positive ion mode by means of an *Autoflex I* mass spectrometer (Bruker). The instrument was controlled by the FlexControl software and equipped with a nitrogen laser (*λ* = 337 nm) operating at a pulse rate of 10 Hz. Pulse ion extraction time was 10 ns and the sampling rate was set to 2.0 gigasamples per second (GS/s). MS measurements were carried out using an acceleration voltage of 20.00 (ion source 1) and 18.45 (ion source 2) kV. Lens voltage was 6.70 kV. Spectra were stored in the range between m/z 700 and 3500. Bruker’s peptide calibration standard II was employed for external calibration.

### Blinded validation of the MALDI-ToF MS protocol for detection of cereulide producing B. cereus strains

All *B. cereus* strains belonging to the test panel, provided blinded by Vetmeduni Vienna to RKI (for details of strains see table 1), were grown on Caso agar for 24 h at 37°C. Colony material was then processed by the ethanol/FA method (see above), which involved transfer into 300 μL of double-distilled water and adding of 900 μL ethanol. From the 75% ethanol fractions a volume of 1 μL was mixed on target with 1 μL of saturated HCCA solution.

In addition, concentrated extracts were prepared by evaporating the remaining supernatants to dryness in a vacuum concentrator (SpeedVac) followed by resolubilization in 20 μL ACN (100%). Volumes of 1 μL of the concentrated extracts were mixed on target with the HCCA matrix solution (see above) before the measurements. MALDI-ToF mass spectra were obtained in positive reflectron mode using the *Autoflex Speed* instrument from Bruker. Microbial samples were considered as cereulide positive if the intensity of the potassium adduct signal at m/z 1191.6 was at least three times larger than the noise intensity. MALDI-ToF MS test results were then transmitted to the Vienna group for unblinding and for comparison with cereulide toxin data from UPLC-TOF MS profiling (Stark et al., 2013).

### Determination of detection limits by MALDI- and matrix-free laser desorption ionization (LDI) –ToFMS

Cereulide standard (50 μg/mL in ACN, product ID CX20422) was purchased from Chiralix (Nijmegen, Netherlands) and used to prepare a dilution series. Cereulide concentration in the diluted ACN solutions varied between 0.5 ng/mL and 1 μg/mL; from these solutions a volume of 1 μL was either directly deposited onto a ground steel target (LDI) and allowed to air dry, or mixed before sample spotting with an equal volume of a saturated solution of HCCA (MALDI). From each of the solutions 5, or 6 target position spots were prepared while one mass spectrum was collected per spot. The LDI dilution series was prepared twice so at least 11, sometimes 12 LDI spectra were collected and analyzed. Mass spectra from the cereulide standard samples were collected in reflector mode as described above, except that each recorded spectrum was obtained by summing up 500 laser shots fired consecutively at each of 20 predefined shot step positions (large spiral geometry, altogether 10,000 laser shots per target position spot). The objective of this highly standardized procedure was to allow reproducible determination of the cereulide amount.

## 4. Results

### Direct cereulide detection from colony materials prepared for bacterial typing

Biotyping by MALDI-ToF MS has become a widely used method for bacterial identification, using mainly ribosomal subunit proteins as target. In order to investigate whether MALDI-ToF MS biotyping workflows established for bacterial identification can be used for direct and sensitive detection of the emetic toxin, cells of *B. cereus* F4810/72 grown on different solid agars, blood, LB and Caso were examined. Preparation of the microbial cells followed the rules and principles of published protocols used for sample processing in biotyping workflows, such as direct transfer, ethanol/FA and TFA extraction. The obtained reflectron MALDI-ToF mass spectra are shown in figure 1.

**Figure 1.**
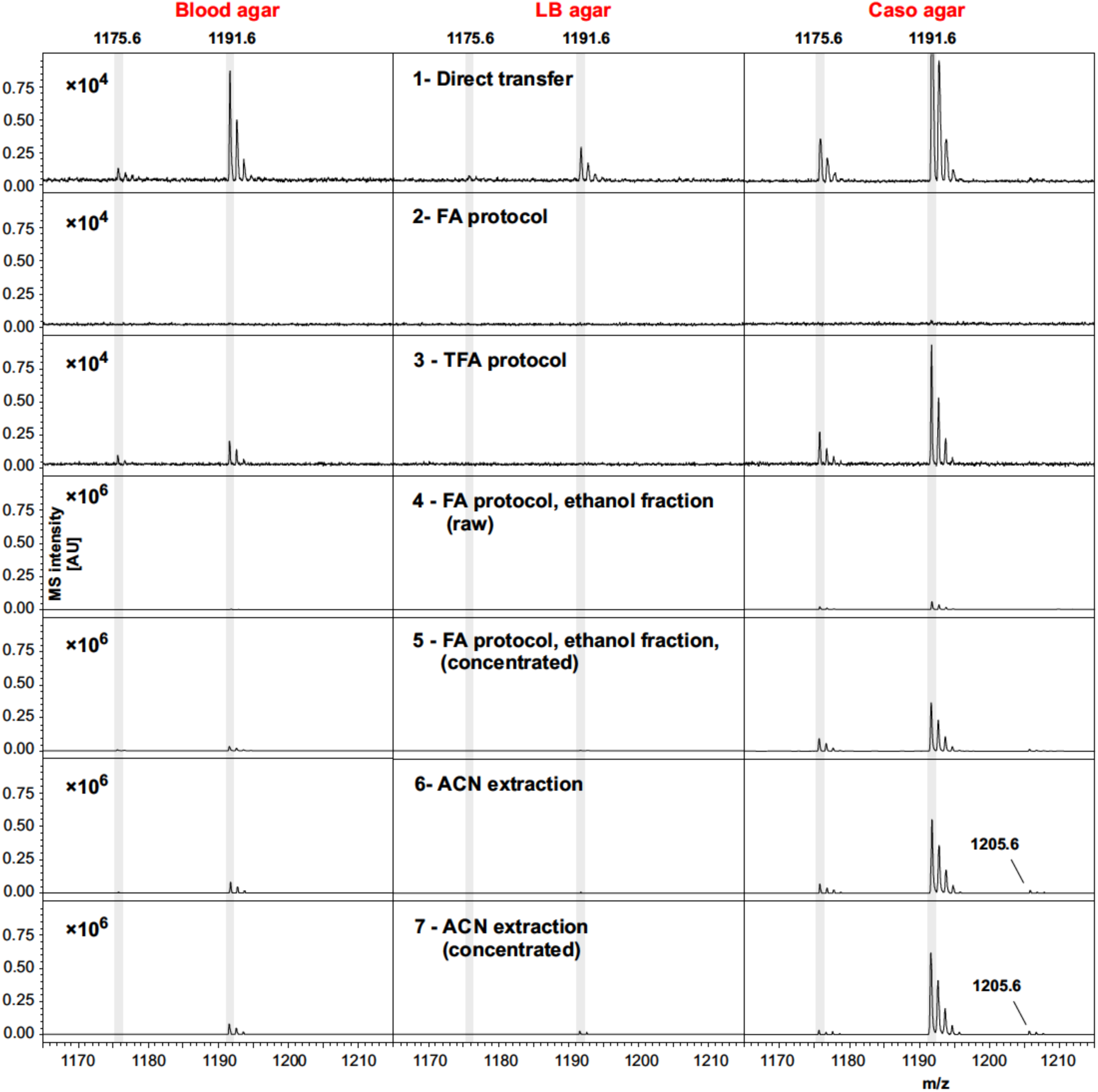
Cereulide detection in *B. cereus* samples cultivated using different cultivation media and different sample preparation, or cereulide extraction methods. Three commonly applied sample preparation techniques for bacterial biotyping workflows, namely the direct transfer (colony-smear) method and ethanol-formic acid (FA), or trifluoroacetic acid (TFA) extraction were utilized (cf. first three rows). In these cases mass spectra were recorded from *B. cereus* F4810/72 samples, or extracts thereof, grown on blood (left column), LB (middle column) or Caso agar (right column) using the positive reflectron mode. Rows 4 and 5 display mass spectra from the ethanol washing solution (raw, or concentrated) obtained when applying the standard ethanol/FA extraction technique. To obtain spectra of row 5, ethanol washing solutions were completely dried and solubilized afterwards in a volume of 20 μL ACN. Spectra from these solutions were recorded after drying 1 μL and adding matrix on target steel plates. Row 6 and 7: Mass spectra acquired after extracting the cereulide by ACN (again as raw and concentrated extracts, respectively). Cereulide was detected as sodium [M+Na]^+^ adduct at m/z 1175.6 and as potassium [M+K]^+^ adduct at m/z 1191.6 (cf. shaded areas). Isocereulide A and/or isocereulide F was detected as potassium [M+K]^+^ adduct at m/z 1205.6 of *B. cereus* samples cultivated on Caso agar (visible only in rows 5-7). Note the different scaling of the intensity axes in rows 1-3 (factor 10^4^) and lines 4-7 (×10^6^). Sample preparation and MS measurements were carried under highly standardized conditions: same matrix, same instrument, 6000 laser shots co-added per sample, nearly identical laser intensity, etc..

In figure 1 mass spectra obtained from *B. cereus* F4810/72 cultures grown on solid blood agar are depicted in the left column whereas the middle and right columns display spectra from LB and Caso agar cultures. Rows 1-3 illustrate reflector mode MALDI-ToF spectra from samples produced by the different preparation methods for MALDI biotyping in the m/z region between 1165 and 1215 (cf. inset of figure 1). Lines 4-5 depict spectra from the washing fraction used in the widely used ethanol/FA extraction protocol, namely the unprocessed (raw) ethanol washing fraction containing 25% water and 75% ethanol (row 4) and the concentrated ethanol fraction (row 5). Note that the intensity scaling in these panels differs from the scaling in rows 1-3 by a factor of 100. Cereulide was detected as sodium [M+Na]^+^ and potassium [M+K]^+^ adduct at m/z 1175.6 and 1191.6, respectively (see shaded areas). The highest intensities of emetic toxin peaks were generally found in preparations, or extracts of Caso agar cultures. Moderate or even low intensities of the peaks at m/z 1175.6 and 1191.6 were typically present in spectra from *B. cereus* cultures grown on blood or LB agar, respectively.

The absence of cereulide signals in samples processed by the ethanol/ FA method might be explained by the wash steps with 75% ethanol solution (see row 2). Indeed, spectra from the raw and concentrated ethanol fractions exhibit mass signals of the cereulide (cf. rows 4 and 5, *B. cereus* cultures cultivated on Caso or blood agar). Thus, cereulide detection from cell extracts obtained by the widely used ethanol/FA extraction method cannot be recommended. Instead, the ethanol washing solution should be used to test for the presence of cereulide.

The effectiveness of cereulide extraction from the microbial samples was improved by replacing of the water/ethanol washing solution by an equal volume of pure ACN (see lines 6 and 7 of figure 1.) Under the given experimental conditions (*B. cereus* F4810/72, same growth, measurement and sample treatment conditions, cultivation on Caso or blood agar), direct detection of cereulide was possible from microbial cells washed by pure ACN.

### Extraction of the toxin from B. cereus colony material by different solvents

Five different organic solvents were tested for their efficacy to extract cereulide from *B. cereus* cultures. For this purpose, equal amounts of cell material from *B. cereus* F4810/72 cultivated on blood agar were used for systematically comparing the intensities of MALDI-ToF MS cereulide signals (see figure 2). In order to eliminate effects of the solvents on the matrix crystallization, all extracts were dried completely and resolubilized in ACN before spotting onto the MALDI target plates. The sodium [M+Na]^+^ and potassium [M+K]^+^ adducts of cereulide were detected at m/z 1175.6 and m/z 1191.6 in all samples with high signal-to-noise (SNR) ratios, however at different signal intensities. The protonated form of the cereulide at 1153.6 [M+H]^+^ could not be detected in any of the spectra. The systematic comparison of different solvents for cereulide extraction revealed, that direct acetone extraction of the bacterial cultures resulted in the highest and n-pentane in the lowest cereulide signals intensities. Nevertheless, ACN and ethanol, resulting in the second and third highest intensity, respectively, were chosen as extraction solvents for further experiments because detection of cereulide was possible in both cases by raw extracts, i.e. without further concentration by vacuum centrifugation. In contrast to ACN and ethanol, acetone is generally less suitable for MALDI-TOF MS because it may trigger the formation of large matrix crystals leading to limited reproducibility. ACN results in a more suitable crystallization whereas ethanol is used as a washing solution in a well-established sample processing method for microbial biotyping (see below).

**Figure 2.**
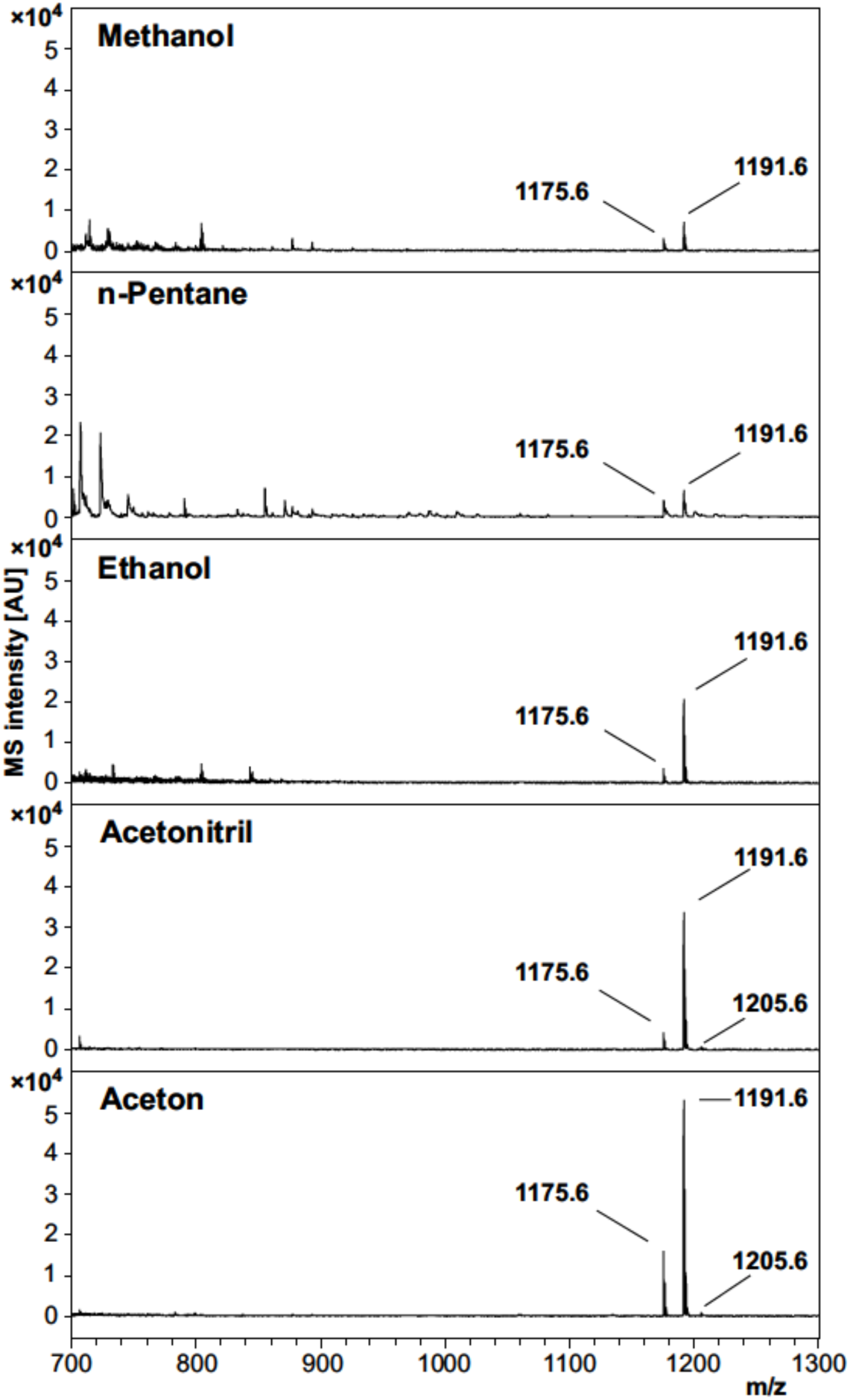
Effectivity of cereulide extraction by different solvents from *B. cereus* F4810/72 colony material. Five solvents, methanol, n-pentane, ethanol, ACN and acetone, were compared for their efficacy to extract the emetic toxin from cells of *B. cereus*. Equal amounts *B. cereus* F4810/72 colony material were used for solvent extraction. MALDI-ToF mass spectra were acquired from concentrated solvent extracts by accumulating altogether 8000 laser shots in the m/z range of 700 – 3500 (reflector mode, 1 μL extract + 1 μL matrix solution). Spectra are equally scaled. Cereulide peaks of each spectrum are labelled with their corresponding m/z value, whereas 1175.6 is the sodium [M+Na]^+^ and 1191.6 the potassium [M+K]^+^ adduct of cereulide. The peak at position 1205.6 is the potassium [M+K]^+^ adduct of isocereulide A and/or isocereulide F. The protonated form [M+H]^+^ of cereulide at m/z 1153.6 could not be detected in any of the mass spectra.

### MALDI LIFT-ToF/ToF MS spectrum of cereulide

In order to confirm the molecular identity of peaks assigned in the literature as sodium ([M+Na]^+^) and potassium ([M+K]^+^) adducts of the cereulide, the peak at m/z 1191.6 has been sequenced by MALDI LIFT ToF/ToF MS. For this purpose, 1 μL of the concentrated ACN extract of *B. cereus* F4810/72 was spotted onto a ground steel target. LIFT-ToF/ToF MS measurements revealed the formation of sodiated and potassiated fragment series (cf. figure 3) of the cereulide. Both fragment series unambiguously confirmed the molecular identity of the peak at m/z 1191.6 as the potassium adduct ion peak of cereulide (see inset of figure 3).

**Figure 3.**
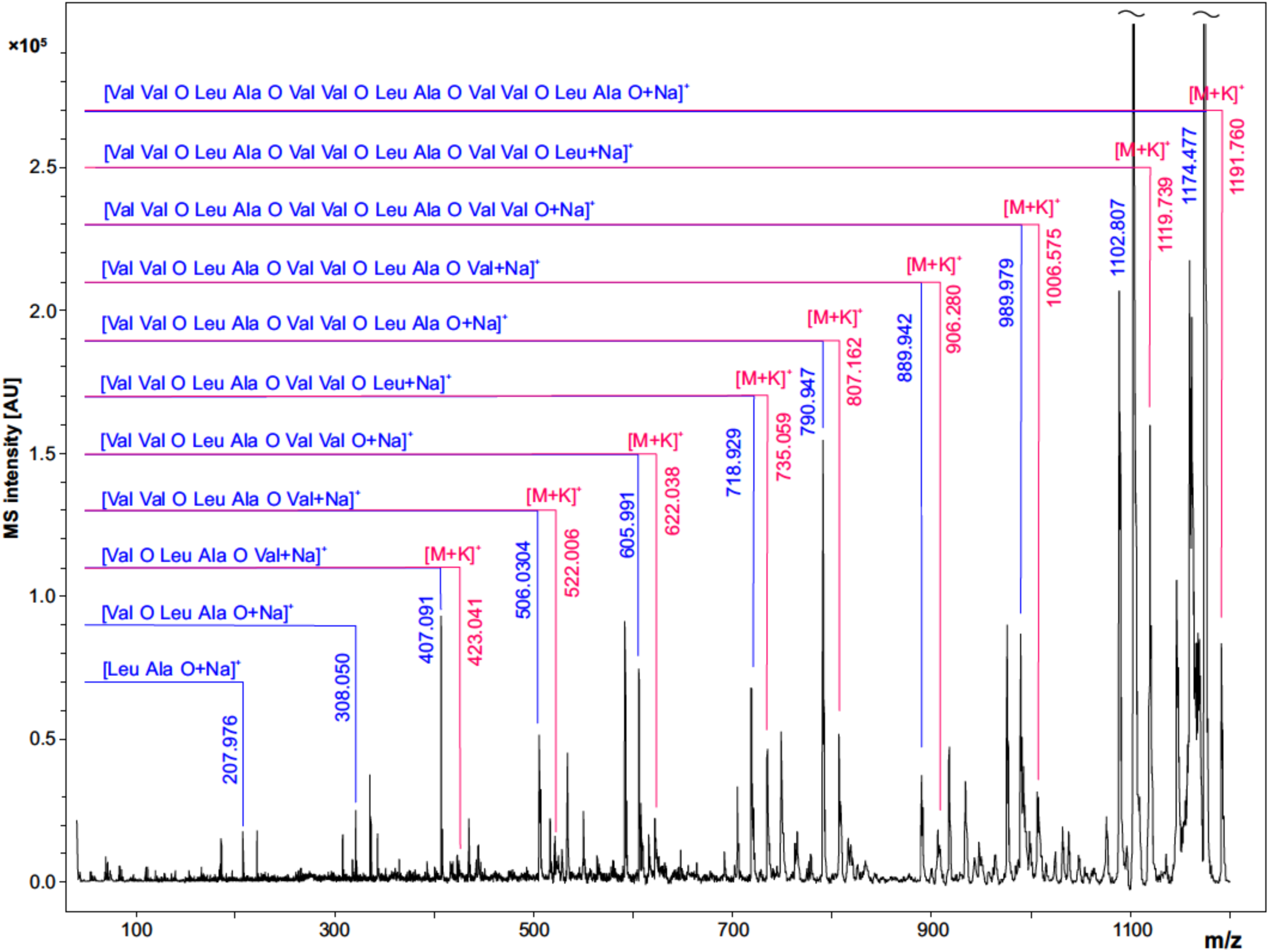
MALDI LIFT-ToF /ToF MS spectrum of cereulide. The potassium [M+K]^+^ adduct of cereulide at m/z 1191.6 was fragmented using laser induced dissociation (LID). Fragmentation resulted in observation of two series of sodiated and potassiated fragment ion adducts in the MALDI LIFT-ToF mass spectrum. The experimental m/z peak positions of these signals enabled assignment to a variety of fragment ions specific for cereulide and thus allowed unambiguous molecular assignment of the toxin.

### Blinded validation of MALDI-ToF MS for cereulide production of B. cereus strains

Cereulide was qualitatively determined by MALDI-ToF MS under blinded conditions from the ethanol washing solutions of a panel of *B. cereus* strains, provided blindly by the Vetmeduni Vienna group to the RKI partner institution (table 1). *B. cereus* strains were selected from the strain collection of the Vetmeduni group to test whether MALDI-ToF MS is suitable for detecting the emetic toxin over a broad concentration range. The test panel comprised a total of eight *B. cereus strains*, including two high-producer, one medium-producer, three low-producer and two none cereulide producing strains. Cereulide production capacity of strains was previously determined by means of stable isotope dilution analysis (SIDA) and HPLC–MS/MS (Stark et al., 2013). The non-emetic *B. cereus* type strain ATCC 14579 and the emetic-like strain ATCC 10987, which is genetically closely related to the emetic *B. cereus* reference strain F4810/72 but is lacking the *ces* genes, were included as negative controls. Spectra were first obtained from raw ethanol washing solutions. In case of negative test results concentrated extracts were prepared and characterized afterwards. All cereulide producing strains were correctly identified using MALDI-ToF MS, as described in the material and method section. Both cereulide negative strains were also correctly classified. It turned out non-concentrated ethanol extracts always contained sufficient amounts of the emetic toxin which demonstrates the high analytical sensitivity of the technique. On overview of the results is provided in table 1.

### Determination of the limit of detection (LOD) of cereulide by MALDI- and LDI-ToF MS

It has been stated recently, that the sensitivity of cereulide detection can be improved by a factor of 1000 using matrix-free laser-desorption/ionization (LDI) MS (Ducrest et al., 2019). However, neither spectra, nor other LDI-derived data such as calibration curves were presented. Therefore, one of the goals of the present study was to investigate the applicability of LDI-ToF MS and to systematically compare the LODs of MALDI- and LDI-ToF MS-based cereulide detection. To this end, a dilution series of a commercial cereulide standard within the range of 0.5 ng/mL to 1 μg/mL was analyzed by performing 5 (MALDI) or 11 (LDI) replicate measurements using standardized MS parameters for data acquisition. The results of these reflectron-mode investigations are illustrated in figure 4. The mean, minimum and maximum intensity values of the potassium peak of cereulide (m/z 1191.6) are plotted against the amount of cereulide per sample spot for MALDI and LDI-ToF MS measurements, respectively. The double-logarithmic representation of the upper panel illustrates successful detection of an amount of 0.25 ng cereulide per sample spot in 3 of 5 MALDI-ToF MS measurements. These data suggest that the LOD under the given experimental conditions (MALDI-ToF MS, reflectron mode measurements, preparations of the pure toxin, *Autoflex Speed* mass spectrometer) equals roughly 0.25 ng (cf. arrow in the top panel of figure 4). Furthermore, despite our large efforts to standardize data acquisition, application of LDI-TOF MS resulted in substantial variations of cereulide signal intensities, which significantly hampered accurate determination of the LOD. The intensity variations of the cereulide’s potassium adduct peak in MALDI- and LDI-ToF MS measurements are illustrated by the blue shaded areas in the upper and lower panel of figure 4. In the LDI case, a cereulide amount of 0.05 ng per spot was sufficient in 11 out of 11 cases whereas [M+K]^+^ peaks of 0.01 ng cereulide were found in 3 out of 11 spectra. These experimental data suggest that LDI-ToF MS improves the detection limit for cereulide by a factor of about 5-25. The improvement of the LOD by such a factor is remarkable, but clearly deviates from the claimed 1000-fold improvement by LDI (Ducrest et al., 2019). Of note are also the technical difficulties linked to LDI measurements. As an illustration, visual assessment of matrix crystals enables an experienced operator to identify suitable measurement positions (*so called* sweet spots) and to select positions with homogeneous crystal structures for subsequent data acquisition. Mass spectra obtained in this operator-guided mode are usually more reproducible and of a higher spectral quality compared to automated measurements. In case of LDI-ToF MS measurements, however, matrix and matrix crystals are absent so that the quality of spectra acquired in the operator-guided mode may vary greatly. In frame of this study, MALDI- and LDI-ToF MS measurements were thus carried out by means of a pre-defined geometry of laser shot step positions (see above). The results of these measurements clearly indicate a low level of reproducibility, which may considered a limiting factor for the applicability of LDI-ToF MS in a routine environment.

**Figure 4.**
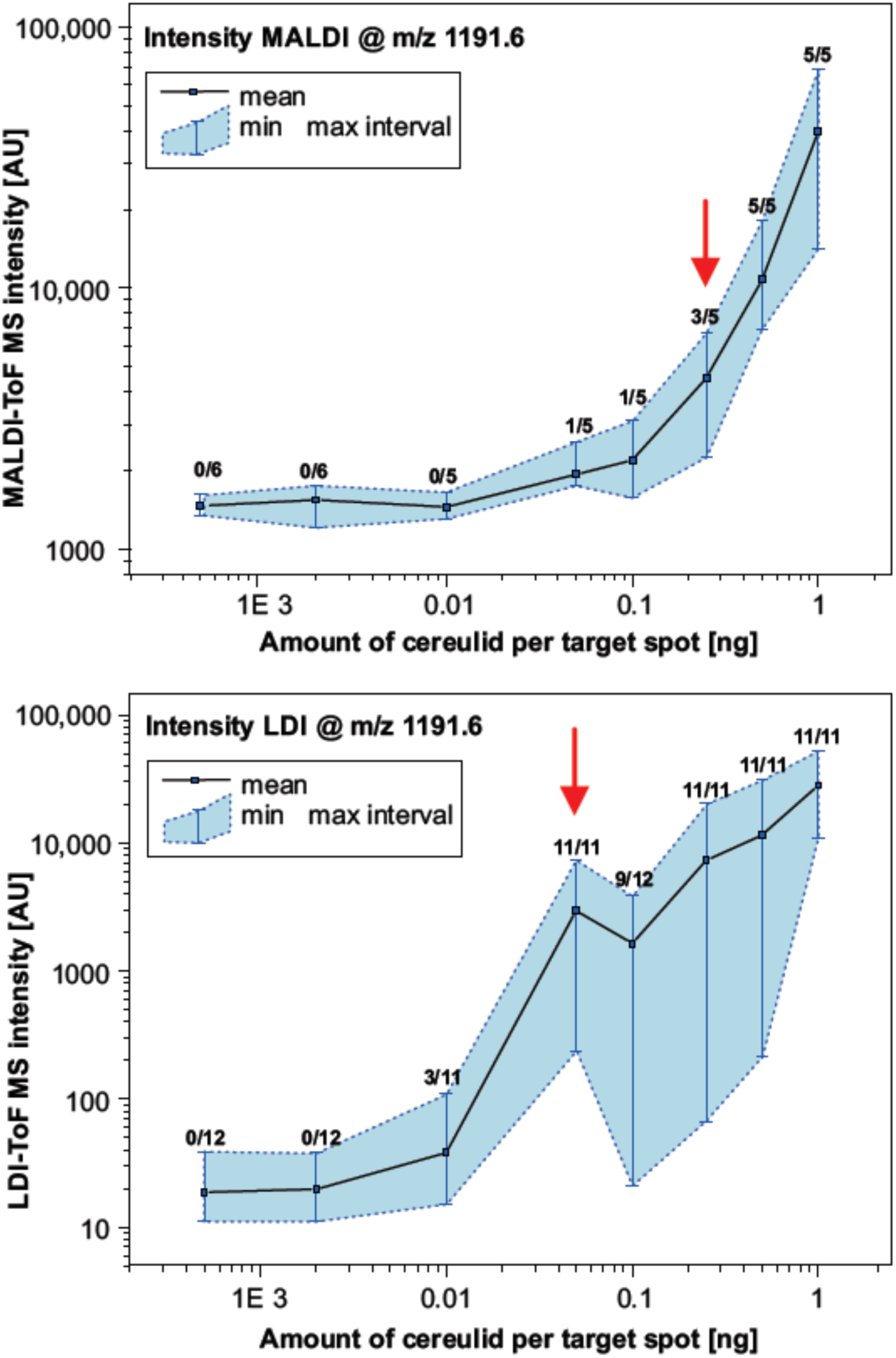
Determination of the limit of detection (LOD) of cereulide by MALDI- and LDI-ToF MS. MALDI-ToF MS (upper panel) and matrix-free LDI-ToF MS (lower panel) were used to determine the LOD of cereulide. For this purpose, a commercial cereulide standard was obtained and used to prepare a dilution series. Mass spectra were collected in reflector mode under strictly standardized conditions. Criteria for a positive test result are (i) a mass peak of the potassium adduct [M+K]^+^ of cereulide at m/z of 1191.6 and (ii) a SNR threshold of at least 3. Data points represent the mean and the min/max MS intensity values from 5 (6) (MALDI) or 11 (12) (LDI) measurements at each point of the dilution series. Furthermore, numbers of positive test results in relation to the total number of replicate measurements are indicated for each point of the curves. Note the double logarithmic scaling (see also text for details).

Selected MALDI- and LDI-ToF MS spectra of the dilution series are presented in figure SI-01 (see supporting information).

### Detection of cereulide in the linear mode of MALDI- and LDI-ToF MS

Bacterial identification is usually performed in an automated fashion using specialized MALDI-ToF MS instruments with dedicated and simplified hardware compared to the one’s utilized for cereulide detection by us at RKI and others (Ducrest et al., 2019;Ulrich et al., 2019). MS instruments for microbial identification often allow m/z determination in linear mode only and offer moderate levels of resolution and mass accuracy compared to reflector-based MALDI-ToF/ToF instruments. Therefore, we performed the analysis in the linear mode also by an instrument designed in the 1990s that is equipped with a low-cost N_2_ laser (*Autoflex I*) to examine if the detection of both cereulide adduct ions is possible by instrumentation similar to mass spectrometers used for microbial biotyping. Selected spectra for MALDI- and LDI-ToF MS are opposed to the respective reflector-based data in figure SI-02. It was found, that cereulide detection is possible in linear mode and that LDI-type measurements improve the peak resolution compared to MALDI. However, as expected, spectral resolution and mass accuracy were lower in linear-mode mass spectra and particularly low in spectra obtained by the *Autoflex* I instrument.

## 5. Discussion

*Bacillus cereus* is a major concern in food production processes because of its high prevalence in food samples related to poisoning (Messelhausser et al., 2014). Due to the high biological activity of the emetic toxin cereulide and the severity of the diseases linked to this highly potent toxin, the detection and identification of emetic strains is of special importance in food microbiology as well as in clinical microbiology diagnostics (Ehling-Schulz and Messelhausser, 2013;Rouzeau-Szynalski et al., 2020). Because of its ease of use, widespread availability of required equipment and its high-throughput potential MALDI-ToF MS is widely used in clinical microbiology for bacterial species identification and would be thus an attractive method for cereulide detection and for discrimination between emetic and non-emetic strains. Recently, it has been proposed that MALDI-ToF MS might represent a suitable method for identifying cereulide producing strains of *B. cereus* (Ulrich et al., 2019) and for cereulide detection directly from bacterial cultures (Ducrest et al., 2019).

Both recently published studies use the colony-smear method, also known as direct transfer, for sample preparation of *B. cereus* cultures and rather advanced MALDI-ToF MS instrumentation compared to the equipment usually used for characterizing bacterial isolates. However, with regard to analytical sensitivity, specificity and applicability of cereulide detection in routine microbiological diagnostics, these studies left a number of questions unanswered. For example, no evaluation and optimization of sample processing was performed and only limited data was provided that confirms the claimed LOD values. Therefore, our study aimed at identifying optimal sample preparation conditions to maximize the analytical sensitivity and to evaluate the fitting of the MALDI-ToF MS cereulide detection method into current diagnostic workflows. Our results of the comparison of common sample preparation methods for bacterial biotyping and the efficiency tests of different organic solvents for cereulide detection open up new possibilities for cereulide detection based on the well-accepted ethanol/FA sample preparation protocol proposed by Bruker in the commercial MALDI Biotyper workflow (Freiwald and Sauer, 2009;Bruker Daltonic GmbH, 2012). We propose a workflow for *B. cereus* identification and cereulide detection, which is illustrated in figure 5. Bacterial cells are prepared using standard ethanol/formic acid (FA) extraction without discarding the wash fraction. Microbial species are identified using standard spectral database matching of experimental MALDI-ToF MS spectra in the m/z range between 2 and 20 kDa and, if *B. cereus* is identified, the wash fraction can be further analyzed by linear mode MALDI-ToF MS measurements in the m/z region between 700 – 2000 Da. If necessary, the sensitivity of this workflow can be increased concentrating the ethanol wash solution. If peaks at m/z 1175 and 1191 are detected in samples identified as *B. cereus* and if the peak position difference equals 16 Th, these peaks are the sodium and potassium adducts of cereulide and therefore the cells must be emetic *B. cereus* strains. The workflow based on the ethanol/FA protocol was successfully evaluated in a blinded test, involving eight *B. cereus* strains with varying cereulide production capacities. Furthermore, as revealed by our current study, the cereulide containing wash solution can be directly subjected to LC-MS/MS, which offers the possibility for unambiguous cereulide identification, isoform discrimination and accurate quantification (data not shown).

**Figure 5.**
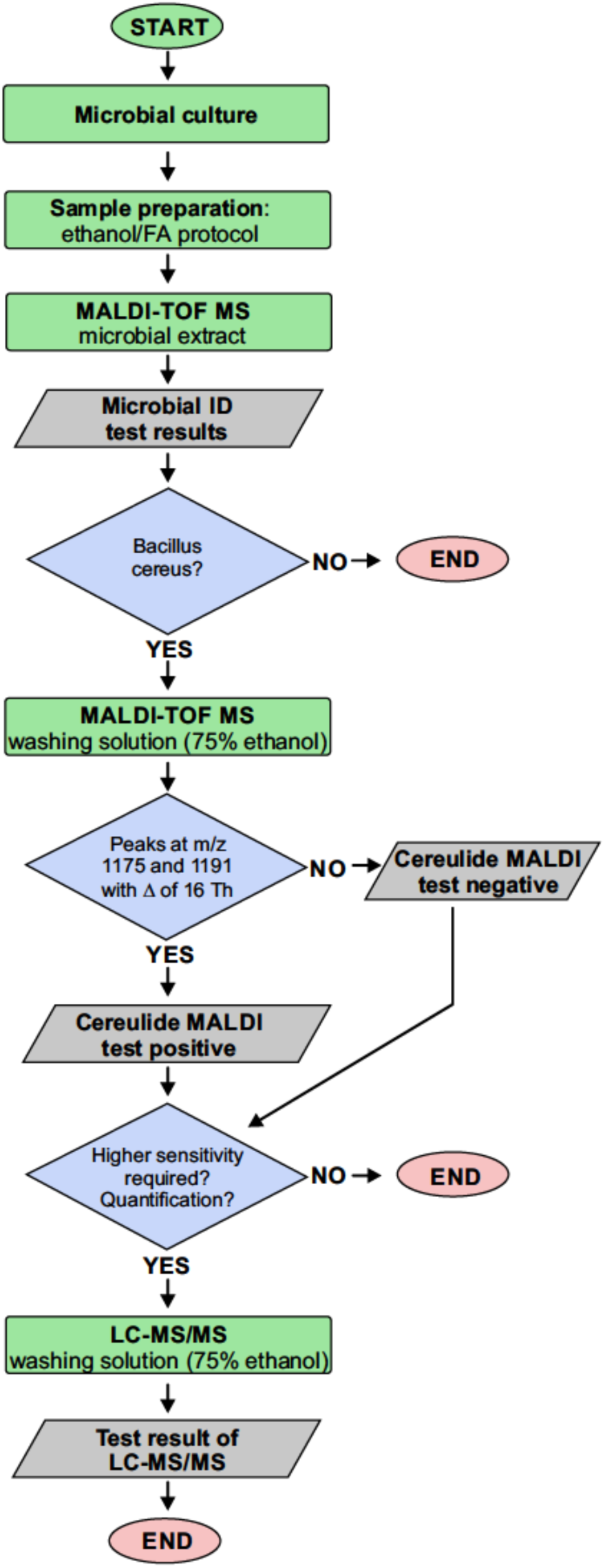
Proposed workflow for cereulide detection based on the ethanol/formic acid (FA) extraction protocol. Bacterial cells are prepared using the ethanol/FA extraction protocol without discarding the wash fraction. Microbial species are identified using standard spectral database matching of experimental MALDI-ToF mass spectra in the m/z range between 2 and 20 kDa. If *B. cereus* is identified, the wash fraction is analyzed in the m/z region between 700 and 2000 Da. The detection of sodium and potassium adduct peaks at m/z 1175 and 1191 with a m/z difference of 16 Th in samples identified as *B. cereus* indicates that these cells are producing cereulide. The wash fraction can furthermore be directly subjected to LC-MS/MS, which offers the possibility for unambiguous cereulide identification, isoform discrimination and accurate quantification.

Selectivity in mass spectrometry is largely dependent on mass accuracy and resolution. However, MALDI biotyping does not require particularly high mass resolution or high mass accuracy. Therefore, instruments specially designed for routine microbial biotyping, such as *Microflex* mass spectrometers from Bruker, are fairly widespread on the one hand but offer only limited options to achieve high mass accuracy and resolution. Such instruments are equipped with a N_2_ laser, a relatively short flight tube and do not allow reflectron mode measurements. Consequently, hitherto rather advanced MS instruments were used for establishing cereulide detection methods (Ducrest et al., 2019;Ulrich et al., 2019). However, since the proposed method should ideally be widely employed to meet the needs of routine diagnostics, we aimed to establish a method for cereulide detection by utilizing entry-level mass spectrometers (linear mode, N_2_ laser, only limited mass resolution, etc.).

Ulrich et al. claimed to have identified two specific biomarkers for cereulide production at m/z 1171 and 1187, which were initially identified by t-tests but later assigned to cereulide itself, because these mass peaks were “similar” to a cereulide standard. Obviously, the biomarker peaks at m/z 1171 and 1187 in Ulrich et al. correspond to the sodium and potassium adduct of cereulide, which were, however, detected in linear positive mode with mass errors of roughly 4000 ppm. This exceeds common mass errors of MALDI-ToF MS for such small molecules by far. The difficulty in the correct peak assignment by Ulrich and colleagues illustrates the specificity problem of MALDI-ToF MS if the molecule identification is solely based on the precursor mass, especially when using linear mode mass spectra of low mass accuracy and resolution. This issue becomes even more apparent when the complexity of the background increases, e.g. if complex samples such as food products are analyzed. Our work and the work of others (Ducrest et al., 2019) showed that fragmentation of cereulide from bacterial isolates using MALDI LIFT-TOF/TOF MS provides sequence information that enables unambiguous toxin detection and enhances selectivity, compared to the assignment of the precursor mass alone (cf. figure 3). However, fragment analysis requires rather advanced MALDI-ToF/ToF MS equipment, usually not available in routine diagnostic laboratories. Nevertheless, the detection of the emetic toxin is possible in linear mode with sufficient certainty (despite the lower resolution and the increased mass error of standard biotyping equipment), as at least two adduct ion peaks are usually detected that must occur at a mass difference of 16 Th. The distance between the sodium and potassium peaks of the cereulide is almost invariant to calibration inaccuracies. In conjunction with bacterial species identification, this approach should enable molecular assignment of mass signals as cereulide at sufficiently high specificity.

Analytical sensitivity is another important aspect of cereulide detection. In the literature, the LOD for cereulide by MALDI-ToF MS was controversially reported as 1 μg/mL (cereulide standard dilution series, linear mode, *Autoflex* Speed (Ulrich et al., 2019)) and 30 ng/mL (cereulide standard, reflectron mode (Ducrest et al., 2019)) or even 30 pg/mL in case of LDI-ToF MS measurements (Ducrest et al., 2019). Of note, the latter data was reported without providing any supporting information, such as spectra, calibration curves, etc. The claimed sensitivity for LDI-ToF MS-based detection of cereulide would correspond to approximately 25 amol cereulide per target spot.

Under our experimental conditions we were unable to confirm the reported massive sensitivity increase by LDI-ToF MS. For reflectron MALDI-ToF MS we determined a LOD for cereulide of 0.25 ng (250 ng/mL) per sample spot which falls in the range between the two reported values. The LOD of LDI-ToF MS was determined to vary between values of 10-50 pg per spot (10-50 ng/mL). This increase of sensitivity was, however, achieved at the costs of technical difficulties and a largely reduced reproducibility (cf. figure 4).

Our proposed workflow for *B. cereus* identification and subsequent cereulide detection (figure 5) was evaluated in a blinded study of high-, medium- and low-producing strains. The proposed workflow, which is based on the ethanol/formic acid (FA) extraction protocol, enables the extraction of cereulide in the ethanol wash fraction from bacterial cultures without any modification of the original FA extraction protocol. All samples were classified correctly using MALDI-ToF MS including one strain which produces only 6.9 μg/mL cereulide. The data revealed that MALDI-ToF MS has a sufficiently low LOD for the analysis of low producing *B. cereus* strains. The cereulide extracts were further verified and quantified by LC-MS/MS. This strategy is a convenient way to obtain retention samples for further cereulide characterization in contrast to the direct transfer (colony-smear) method and fits well into current workflows of microbiological diagnostic laboratories. Another benefit of toxin extraction is the ability to concentrate the ethanol extracts and thereby enhance the sensitivity of cereulide detection.

## 6. Conclusion

The ease of use and the wide-spread availability of MALDI-ToF MS equipment should be helpful to identify cereulide producing *B. cereus* strains in a routine setup and to diagnose food-associated cereulide intoxications by a simple, rapid and cost-effective technology. The proposed workflow is based on the well-established ethanol/formic acid (FA) extraction protocol and can be used without any modification of a sample preparation procedure routinely used for bacterial identification.

## 7. Acknowledgments

The authors wish to thank D. Jacob (RKI, Highly Pathogenic Microorganisms, ZBS 2) for providing emetic and non-emetic test strains of *B. cereus*. Furthermore, the authors are grateful to J. Rau (Chemisches und Veterinäruntersuchungsamt Stuttgart, Germany) for fruitful discussions and support.

## 8. Author Contributions Statement

PL, MES, AS and JD contributed to the conception and design of the study; AS and TS collected the data and performed the experiments; PL, MES, TS and JD analyzed and interpreted the data, JD wrote the first draft of the manuscript; All authors wrote sections of the manuscript, contributed to manuscript revision, read and approved the final version of the manuscript

## Abbreviations

ACN: acetonitrile
FA: formic acid
HCCA: α-cyano-4-hydroxycinnamic acid
LC: liquid-chromatography
LDI: laser desorption/ionization
LOD: limit of detection
MALDI-ToF: matrix-assisted laser desorption/ionization - time–of–flight
MS: mass spectrometry
NRPS: non-ribosomal peptide synthetase
SIDA: stable isotope dilution analysis
SNR: signal-to-noise ratio
Th: Thomson
TFA: trifluoroacetic acid

