## Supporting Information for "Evaluation of MALDI-ToF Mass Spectrometry for Rapid Detection of Cereulide from *Bacillus cereus* Cultures"

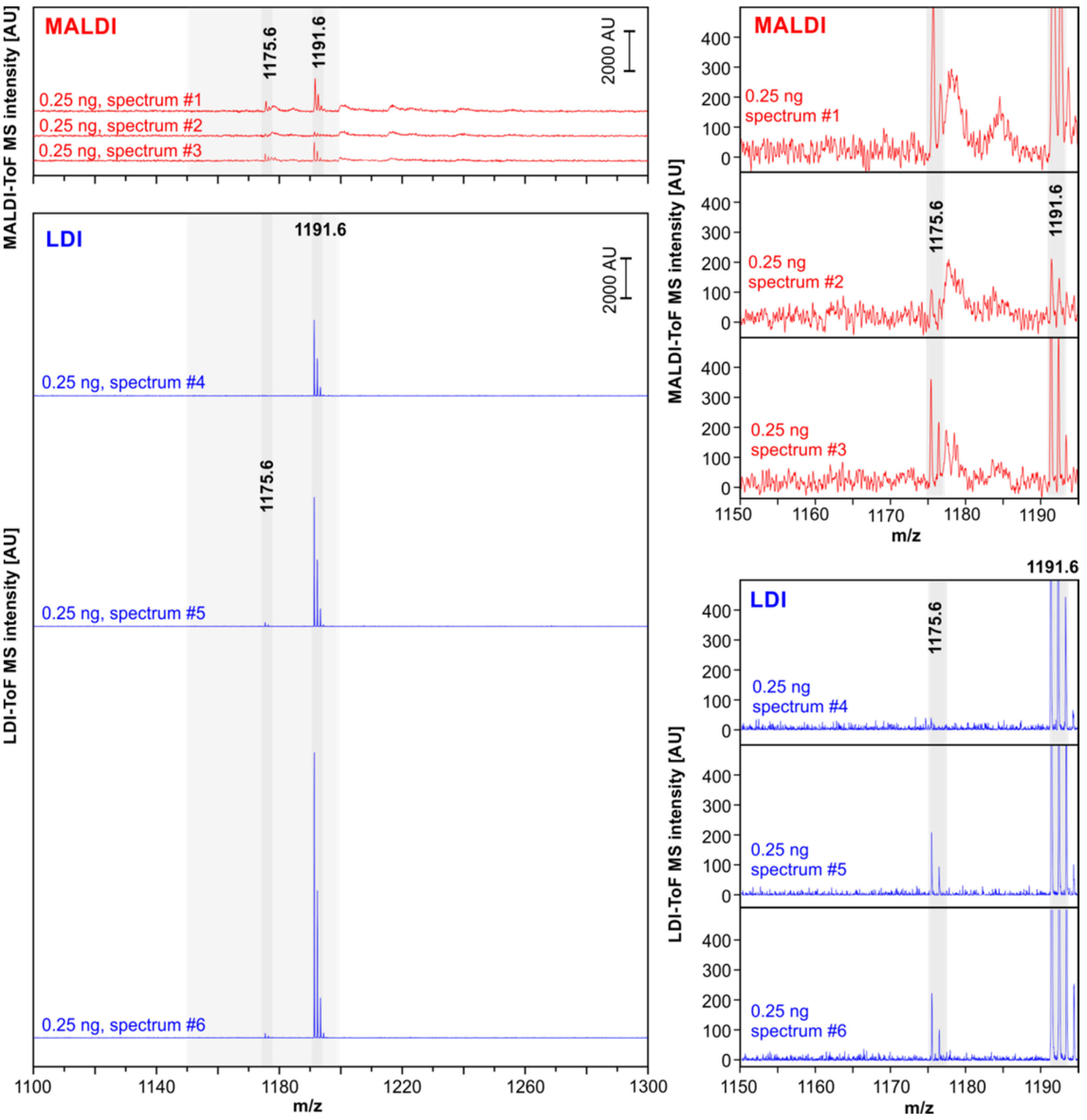


**Figure SI-1.** A selection of MALDI-ToF and LDI-ToF technical replicate mass spectra obtained from a commercial cereulide standard. All mass spectra were obtained in reflectron mode from a dried cereulide preparation which contained 0.25 ng of the emetic toxin per target position spot. MALDI-ToF (red) and LDI-ToF mass spectra (blue) are equally scaled to illustrate the intensity differences of the sodium $[M+Na]^+$ adduct peaks at m/z 1175.6 and the potassium $[M+K]^+$ adduct peaks of cereulide at m/z 1191.6 (see also shaded areas).

*Left column*: mass spectra in the m/z region of 1100 – 1300. Spectra #1-#3 (MALDI, red) and #4-#6 (LDI, blue) were selected from the series of MALDI-ToF and LDI-ToF spectra acquired to determine the LOD of cereulide (cf. figure 4). At 0.25 ng of the emetic toxin per target spot, both types of spectra demonstrate significant intensity variations. Cereulide adduct ion peaks in LDI spectra are generally more intense when low toxin amounts are tested.

*Right panels*: enlarged view of the same MALDI- or LDI-ToF mass spectra shown in the left panels. Again, spectra are equally scaled. Note the higher resolution and lower noise level of LDI-ToF mass spectra.

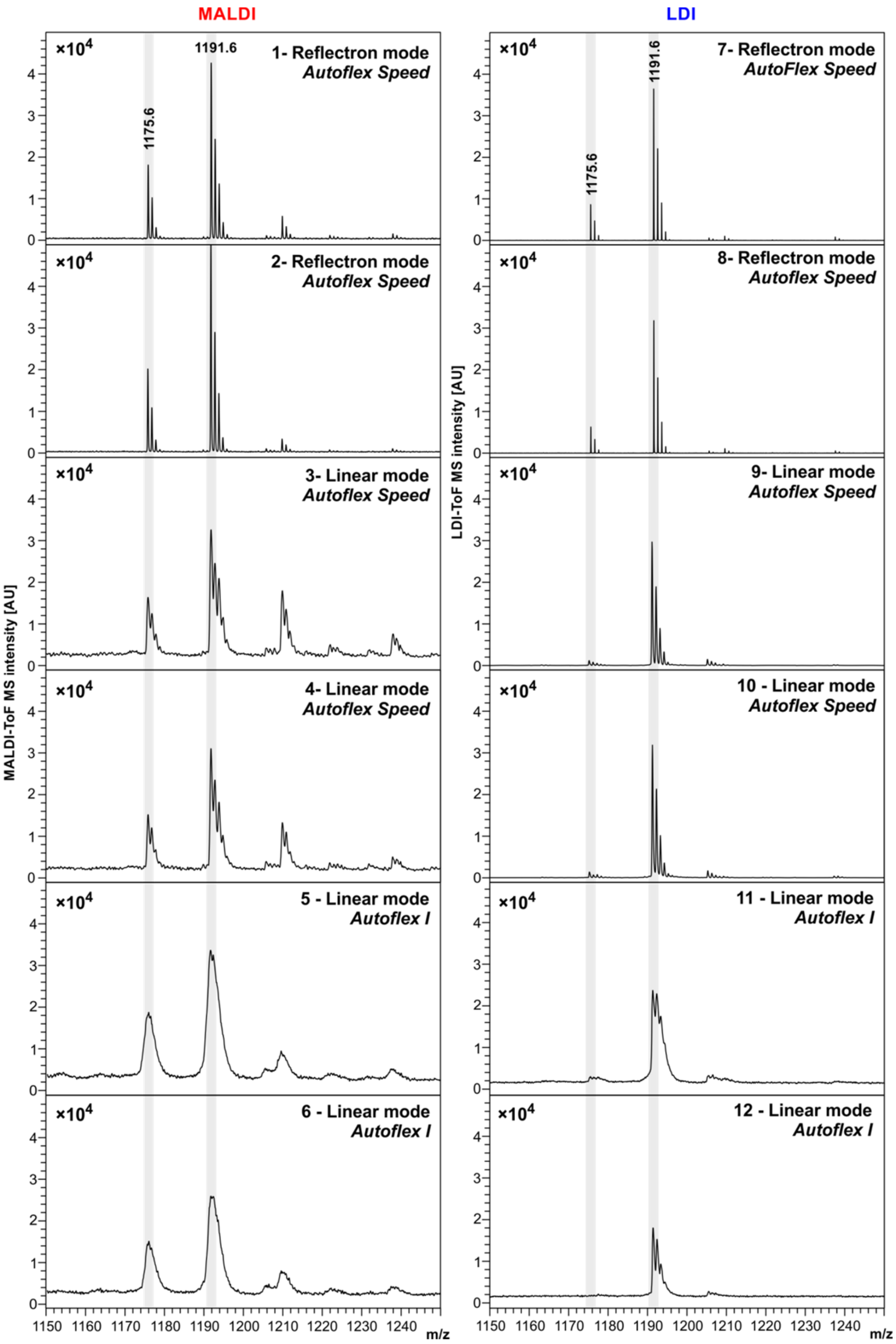
MALDI
LDI
×10^4
1175.6
1191.6
1- Reflectron mode
Autoflex Speed
2- Reflectron mode
Autoflex Speed
3- Linear mode
Autoflex Speed
4- Linear mode
Autoflex Speed
5 - Linear mode
Autoflex I
6 - Linear mode
Autoflex I
7- Reflectron mode
AutoFlex Speed
8- Reflectron mode
Autoflex Speed
9- Linear mode
Autoflex Speed
10 - Linear mode
Autoflex Speed
11 - Linear mode
Autoflex I
12 - Linear mode
Autoflex I
MALDI-ToF MS intensity [AU]
LDI-ToF MS intensity [AU]
m/z
1150 1160 1170 1180 1190 1200 1210 1220 1230 1240

**Figure SI-2.** Direct cereulide detection by means of MALDI- (right, 1-6) and LDI-ToF MS (left, 7-12) in linear and reflectron measurement mode (cf. insets). Samples were prepared by the ethanol/FA extraction method from *B. cereus* F4810/72 cultures grown at 37°C for 24h on Caso agar. 1 µL of the raw ethanol washing solution containing 75% ethanol, non-concentrated) were directly deposited (LDI) or mixed with 1 µL HCCA matrix solution (MALDI) on target and dried subsequently on a stainless-steel target. MS measurements were carried out under highly standardized conditions in the reflectron and linear mode using an *Autoflex Speed* device equipped with sophisticated Smartbeam™ (Nd:YAG) laser technology (panels 1-4 and 7-10), or an *Autoflex I* ($N_2$ laser) mass spectrometer (panels 5-6 and 11-12), both from Bruker. Cereulide peaks are discernible at m/z 1175.6 as sodium adduct $[M+Na]^+$ and at m/z 1191.6 as potassium adduct $[M+K]^+$ of the cereulide. Peaks at m/z 1205.6 represent the potassium $[M+K]^+$ adduct of isocereulide A and/or isocereulide F.

LDI-ToF mass spectra generally exhibit higher resolution and a decreased noise level in both, linear and reflector mode measurements. Furthermore, peaks in linear mode spectra exhibit a reduced mass accuracy and a lower resolution, compared with reflector mode spectra. In addition, the limitations of entry-level equipment in terms of mass accuracy and spectral resolution are illustrated (cf. panels 3 and 4 vs. 5 and 6).
